# An information-theoretic approach for measuring the distance of organ tissue samples using their transcriptomic signatures

**DOI:** 10.1101/2020.01.23.917245

**Authors:** Dimitris V. Manatakis, Aaron VanDevender, Elias S. Manolakos

## Abstract

**Motivation:** Recapitulating aspects of human organ functions using in-vitro (e.g., plates, transwells, etc.), in-vivo (e.g., mouse, rat, etc.), or ex-vivo (e.g., organ chips, 3D systems, etc.) organ models are of paramount importance for precision medicine and drug discovery. It will allow us to identify potential side effects and test the effectiveness of therapeutic approaches early in their design phase and will inform the development of accurate disease models. Developing mathematical methods to reliably compare the “distance/similarity” of organ models from/to the real human organ they represent is an understudied problem with important applications in biomedicine and tissue engineering.

**Results:** We introduce the Transctiptomic Signature Distance, *TSD*, an information-theoretic distance for assessing the transcriptomic similarity of two tissue samples, or two groups of tissue samples. In developing *TSD*, we are leveraging next-generation sequencing data and information retrieved from well-curated databases providing signature gene sets characteristic for human organs. We present the justification and mathematical development of the new distance and demonstrate its effectiveness in different scenarios of practical importance using several publicly available RNA-seq datasets.

**Supplementary information:** Supplementary data are available at *bioRxiv*.

## 1 Introduction

Assessing the transcriptomic distance of biological samples (e.g., organ tissues, cells of different types, etc.) is essential for understanding their functional differences and recognizing disease states (Aibar *et al*. (2016); Crow *et al*. (2019); Mohammed *et al*. (2019); McDonough *et al*. (2019)). Recently, significant efforts have been invested towards characterizing organ tissues (e.g., liver, kidney, intestine, etc.) of different species (e.g., human, mouse, rat, etc.) at various states (e.g., healthy, diseased, etc.) (Uhlen *et al*. (2015, 2017); Yu *et al*. (2015); Keen *et al*. (2015); Mele *et al*. (2015); Suntsova *et al*. (2019); Sollner *et al*. (2017)) using RNA-sequencing, a mature technology for quantifying gene transcripts in biological samples. A notable effort is the Human Protein Atlas (HPA) project (Uhlen *et al*. (2015)), a Swedish-based program providing, among others, gene expression signatures for 37 healthy human organ tissues. In particular, the HPA provides for every tissue type a set of genes exhibiting significantly elevated expressions compared to the other organ tissue types. It is widely accepted that these gene sets form a “transcriptomic signature” of the specific organ, and their expression patterns characterize the tissue’s underlying biological processes.

Recent advancements in bioengineering and biotechnology have enabled the development of cell-cultured based organ models that recapitulate critical functions of human organs (e.g., liver, intestine, brain. etc.) (Jang *et al*. (2019); Kasendra *et al*. (2019)). The emerge of such ex-vivo human organ models generated in turn the need for new mathematical tools for determining their “similarity” to the actual human organ they represent. Such tools will not only help us understand the model capabilities and limitations but also reveal aspects we can improve in their design to optimize their physiological relevance and increase their value for precision medicine. Next-generation sequencing data (e.g., RNA-seq) is already utilized to determine the distance between organ tissue samples (Mele *et al*. (2015); Suntsova *et al*. (2019); Sollner *et al*. (2017); Sudmant *et al*. (2015)) in conjunction with classical mathematical tools such as the Euclidean distance, or dimensionality reduction techniques (e.g., Principal Component Analysis, Linear Discriminant Analysis, Uniform Manifold Approximation and Projection, etc.). However, all these methods exhibit significant limitations due to sensitivity to noisy measurements and outliers, inability to capture non-linear relations, etc. (Pereira *et al*. (2009); Li *et al*. (2016)). In this work, we introduce a new distance, called *Transcriptomic Signature Distance (TSD)*, that was inspired from the field of information retrieval, addresses the problem of tissue sample comparisons in the well-established framework of information theory, and circumvents the above-mentioned weaknesses of more classical approaches.

Text similarity is a well-studied problem in information retrieval (Pradhan *et al*. (2015); Nagwani *et al*. (2015)). Over the years, many techniques have been proposed to measure the distance/similarity of documents based on features such as word frequencies, word patterns in sentences, etc. They process vector representations of documents and assume that documents with similar content exhibit similar feature patterns. RNA sequencing, on the other hand, allows us to read the transcriptome (i.e., read the stories) of tissues. These transcriptome “stories” are written using a four nucleotide bases alphabet, which is applied to construct words (i.e., the genes). Based on this analogy, the transcriptome of homologous tissues should “tell” similar stories, and therefore the set of words (i.e., genes), their relative frequencies of appearance, and rankings in the stories are expected to be similar as well.

Our method exploits well-curated databases (e.g., HPA, GTExPortal, etc.) to retrieve genes that are considered transcriptomic signatures of the compared organ tissues (Yu *et al*. (2015); Lonsdale *et al*. (2013)). These are sizeable gene sets that can adequately characterize the “identity” of organ tissues. Using them as the basis in comparing tissue samples allows us to significantly reduce the effects of sequencing “background noise” and donor-to-donor variability in the analysis. Moreover, making use of information theory and advanced statistical methods, *TSD* can capture the distance/similarity between any two arbitrary organ tissue samples, or between two tissue samples that are known to belong to two different classes (sample groups, e.g., different organs, organ models etc.). In the latter case, *TSD* considers, in a principled manner, the intra-class variabilities and incorporates them into the distance estimation. The proposed distance space is determined using the expressions probability distribution of the signature genes as well as their rankings profile in transcriptomes of the two tissues. We explain the advantages of the proposed distance and experimentally validate its ability to resolve transcriptomic distances between organ tissues in many practical situations of interest.

The rest of the paper is organized as follows: In Section 2, we present the development of *TSD* and justify why it can better capture the actual transcriptomic distance between two tissues. In Section 3, we present and discuss extensive experimental validation results in different scenarios of practical importance. Finally, we summarize our findings in Section 4.

## 2 Methods

In this section, we present *TSD* a new method to measure the transcriptomic distance between organ tissue samples. *TSD* requires as input the gene expression levels (e.g., hit-counts, CPM, TPM, FPKM, etc.) of two tissue samples (or two sets of tissue samples) where one tissue sample is assumed to be the “reference” (i.e., gold standard) from which we want to measure the distance of the other tissue sample.

### 2.1 Preliminaries

We use lowercase letters to denote scalars, bold lowercase (uppercase) letters for vectors (matrices), and bold uppercase calligraphic letters for sets.

Let 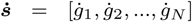 and 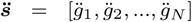 be two vectors storing the expression levels of *N* genes after applying RNA-sequencing on tissue samples 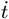 and 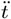 respectively. For presentation purposes we assume that 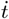 is our *“reference”* tissue sample and 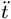 the sample of tissue that we want to measure its distance from 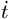. From the Human Protein Atlas database we retrieve the *M* ≤ *N* genes that characterize the reference organ where tissue 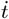 was sampled from and are a subset of genes {*g*_1_, *g*_2_, …, *g*_*N*_}. Then, using 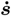 and 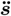 we form the corresponding Atlas signature vectors 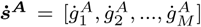 and 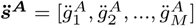. For each Atlas signature vector, we estimate the corresponding discrete probability distribution 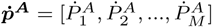 and 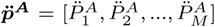. The probabilities of each Atlas gene are calculated using:

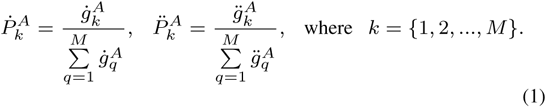

In addition, we form the vectors 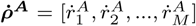 and 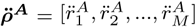 containing the expression level rankings of the Atlas genes of the reference tissue 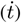 in the full transcriptome gene expression vectors 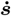 and 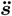. Note that 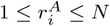.

In Section 2.2, we present the “simple” version of the *TSD* that measures the transcriptomic distance between any two organ tissue samples. In Section 2.3 we present the development of the more general version, the so called *weighted-TSD (wTSD)*, used to estimate the distance between two samples knowing that they belong to two different classes (tissue sets) and taking into account the intra class variabilities.

### 2.2 The Transcriptomic Signature Distance

The proposed *TSD* measures the transcriptomic distance between a tissue sample 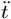 from a *“reference”* tissue sample 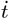 as the average of the *Jensen-Shannon Divergence (JSD)* (Section 2.2.1) and the *Rankings Correlation Distance (RCD)* (Section 2.2.2). We present below the two distances and their limitations when used in isolation that justifies their combined use in assessing tissue distances.

#### 2.2.1 The Jensen Shannon Divergence

*JSD* is popular for measuring the similarity between two probability distributions and is related to Shannon’s entropy, *Kullback-Leibler Divergence (KLD)* and mutual information (Fuglede *et al*. (2004)). *JSD* calculates the divergence (distance) between the probability distributions 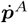 and 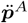 using the following equation:

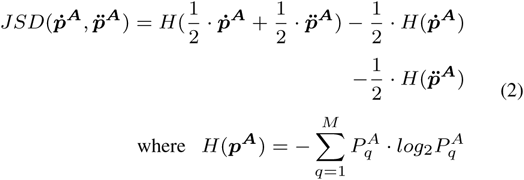

is the Shannon’s entropy function (Jianhua *et al*. (1991)). Unlike *KLD*, the square root of the *JSD* (*SR-JSD*) satisfies all the basic properties of a “true” metric, such as symmetry, non-negativity, triangle inequality and identity of indiscernibles. Given that we use the base-2 logarithm for calculating the Shannon’s entropy (see equation (2)), the *SR-JSD* is bounded in the interval [0, 1] where 0(1) corresponds to minimum (maximum) distance.

*SR-JSD* uses the probability distributions of the reference Atlas genes to measure the distance between the two samples and therefore it does not take into account the expression levels of these genes in the whole transcriptome which is very important for the development of an accurate tissue distance metric. The following example demonstrates this limitation.

Figure 1a shows the whole gene expression profiles (*N* = 20 w.l.o.g) of two organ tissues 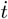 and 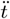. Let’s assume that 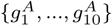 are the reference Atlas genes (*M* = 10) that characterize tissue 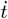. Using their expressions in both tissues (see Figure 1b), we form vectors 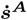 and 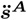 and calculate the corresponding probability distributions 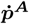 and 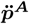 (Figure 1c). Since both tissues in this example 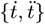 exhibit proportional gene expression for the Atlas genes, this results to identical probability distributions (Figure 1c) and therefore 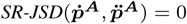, suggesting that 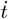 and 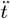 are transctiptomically very close. However, as we can see in Figure 1a this is apparently not the case, which shows that using *SR-JSD* alone may fail to capture tissues transcriptomic distance. To deal with this limitation, we propose to also employ in TSD the correlation coefficient of the rankings of the Atlas genes in the whole transcriptome that can correct this situation (see Figures 1d and 1e).

**Fig. 1.**
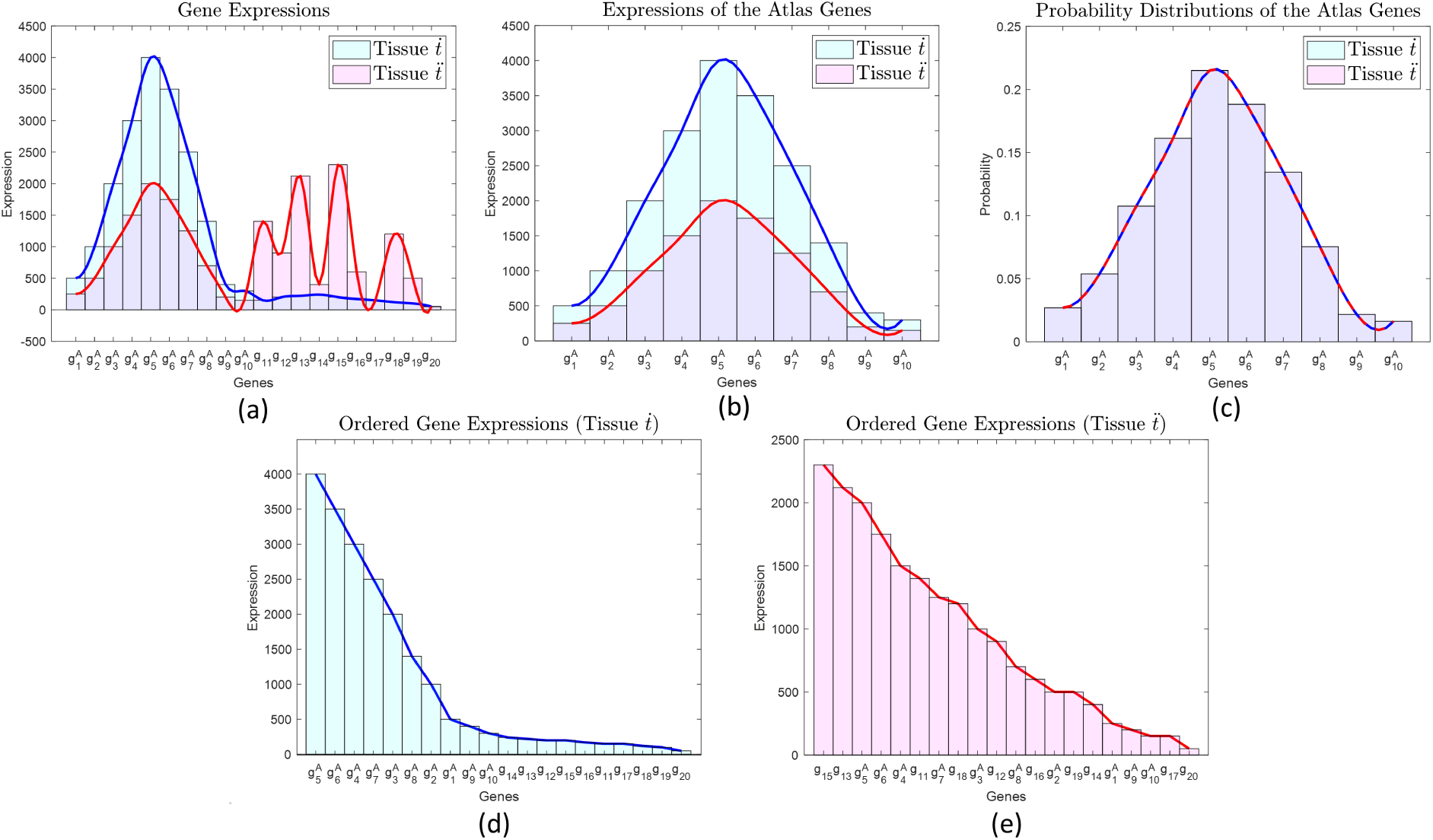
(a) Gene expression vectors of the two tissues 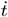 (reference) and 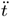. (b) Expression histograms of the Atlas genes 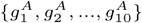 of the tissues 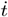 (reference) and 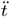. (c) The discrete probability distributions 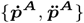 of the Atlas genes are identical due to the proportional gene expression levels (see Figure1b) and therefore 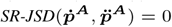. (d)-(e) The gene expression rankings of the tissues 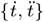. Note that the rankings of the Atlas genes in the whole transcriptome 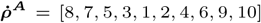 and 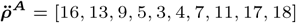 respectively differ significantly which captures the tissue differences in this case.

#### 2.2.2 The Rankings Correlation Coefficient

The Rankings Correlation Coefficient (RCC) is defined as the Pearson’s correlation of the Atlas gene ranking vectors, namely 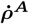 and 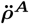, in the whole transcriptome. It is calculated based on the formula:

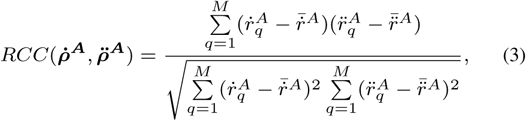

where 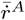 and 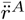 are the corresponding mean rankings. The range of RCC is [-1, 1] where 1(−1) implies perfect linear relation between the compared ranking vectors.

Applying equation (3) to the Atlas ranking vectors provides us information about the linear relation of the reference Atlas genes in the whole transcriptome of the two tissues based on their expressions. Figures 1d and 1e demonstrate the ranking profile difference of the Atlas genes in the transcriptomes of 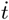 and 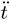 respectively. This difference can successfully capture the tissues dissimilarity when *SR-JSD* may fail to do so as in the example of Figure 1.

#### 2.2.3 Transcriptomic Signature Distance

The example presented in Figure 1 demonstrates the limitation of the *SR-JSD* to measure with accuracy the transcriptomic signature distance between two tissues. A similar example that demonstrates a corresponding limitation when *RCC* is used alone, is provided in Figure 2. In this example, tissues 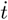 and 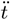 have identical reference Atlas gene rankings in the corresponding transcriptomes (Figure 2d and 2e) but different probability distributions (Figure 2c). In this case, using *RCC* alone would suggest that the transcriptomic signatures of the tissues are identical which is not the case. To address the limitations introduced when using either *SR-JSD* or *RCC* independently, we introduce the *Transcriptomic Signature Distance (TSD*) which combines them:

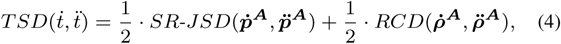

where 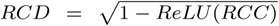 is the *Rankings Correlation Distance. ReLU* is the *Rectified Linear Unit* activation function defined as: *ReLU* (*x*) = *x*, when *x >* 0 and zero otherwise. For the calculation of the *RCD* we assume that if two ranking vectors have *RCC <* 0 (i.e. are anti-correlated) then their Rankings Correlation Distance is maximal (i.e. 1). Note that *TSD* is bounded in the interval [0, 1] where 0(1) corresponds to minimum (maximum) distance.

**Fig. 2.**
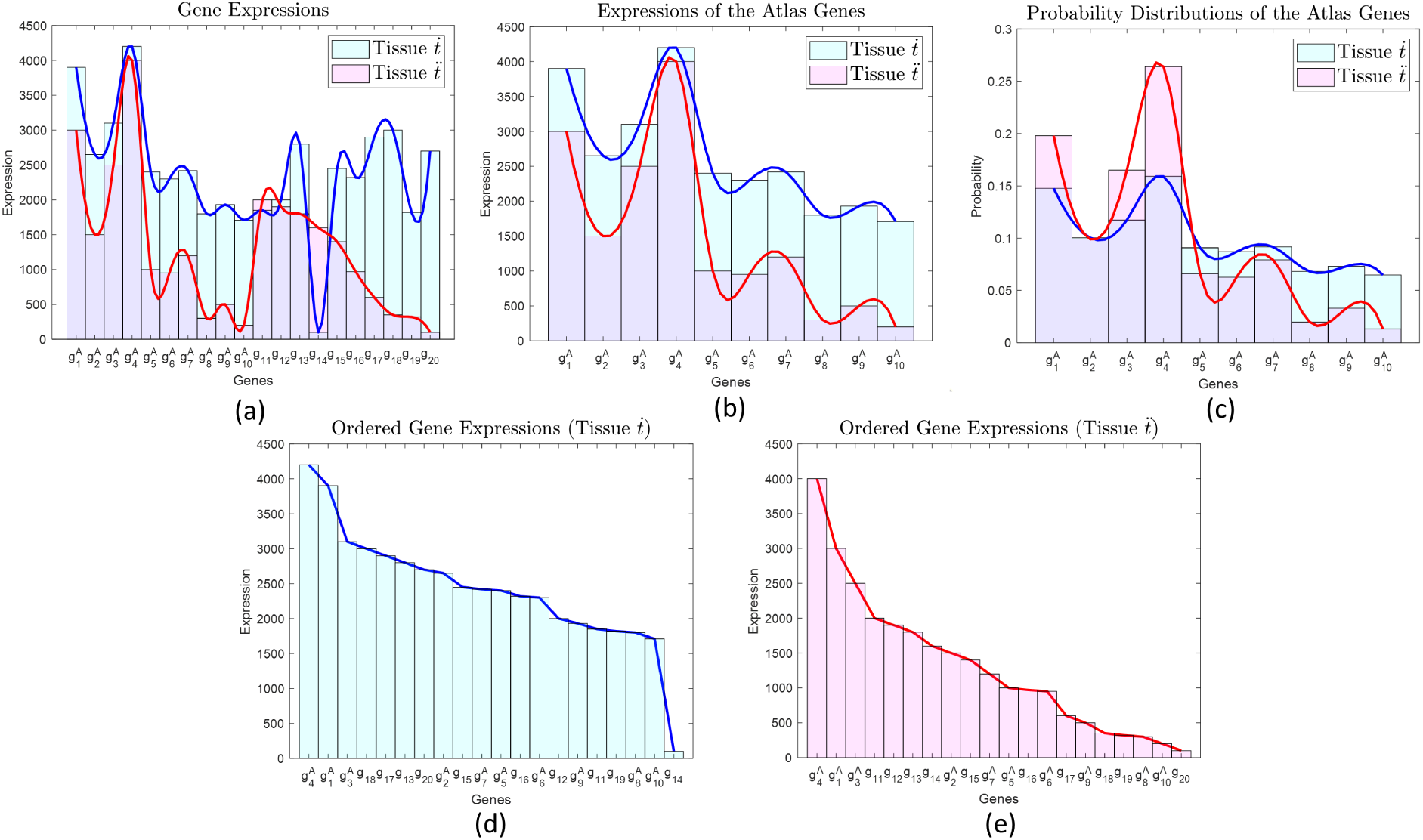
(a) Gene expression vectors of two tissues 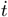 (reference) and 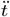. (b) Expression histograms of the Atlas genes 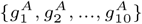 of the tissues 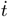 (reference) and 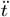. (c) The discrete probability distributions 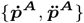 of the reference Atlas genes can capture the gene expression differences of the tissues. (d)-(e) The sorted gene expression profiles of the two tissues. Note that the rankings 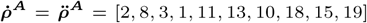 of the Atlas genes in the transcriptome are identical for both tissues, which results to *RCC* = 1. In this example, using the RCC of the Atlas genes alone fails to capture the transcriptomic differences of the two tissues.

### 2.3 Transcriptomic Signature Distance of samples belonging to two different tissue sets

In Section 2.2 we presented *TSD* that can measure the distance between any two tissue samples 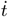 and 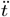 without considering their classification. Here we study the case where we want to measure the distance between two samples knowing that they belong to two different tissue sets (e.g., ex-vivo organ model samples vs. human organ samples). We propose a modified version of the *TSD*, which takes into account the gene expression variability of the samples within the corresponding tissue sets and provides statistically robust and accurate estimations of their transcriptomic signature distances.

Let’s assume that we have two sets of tissue samples 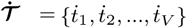 and 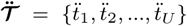. Similarly to the notation used in Section 2.1, 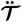 corresponds to the set of tissue samples that we want to compare to the reference set 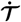. For each tissue sample 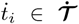, where *i* = {1, 2, …, *V*}, and 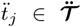, where *j* = {1, 2, …, *U*}, we form the: (i) gene expression profile vectors 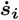 and 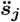; (ii) the Atlas signature vectors 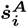 and 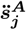; (iii) the discrete probability distributions of the Atlas genes 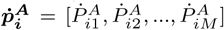 and 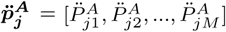; and (iv) the matrices 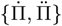 that summarize the Atlas gene probability distributions of the corresponding samples.

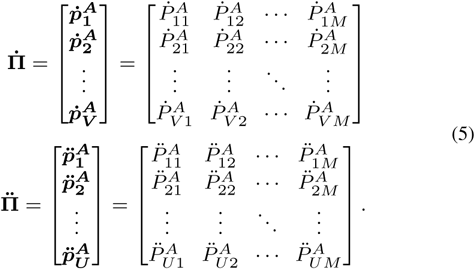

#### 2.3.1 The weighted Jensen-Shannon Divergence

In 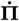 and 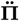, we assume that the probabilities of appearance of each Atlas gene (e.g. *k*^*th*^ gene) across tissue samples (i.e. rows of matrices), were generated by a normal distribution (e.g. 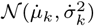 and 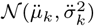) with parameters:

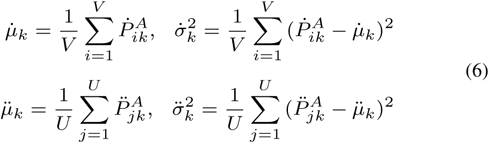

Using this assumption, for each tissue 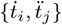, the likelihood of appearance of the *k*^*th*^ Atlas gene can be calculated as:

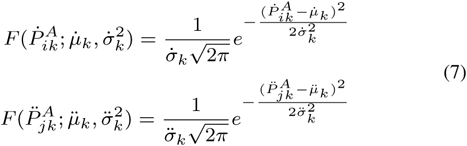

The larger the 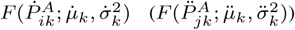 the more “confident” we are about the likelihood of appearance of the *k*^*th*^ Atlas gene in the set of samples 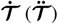. We quantify our “confidence” as:

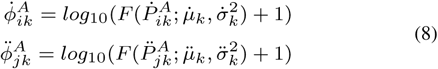

where to avoid negative “confidence” values we added “1” before taking the logarithm of the likelihoods.

To incorporate our “confidence” about the likelihood of appearance of the Atlas genes in the *JSD* we utilize a weighted version of the Shannon’s entropy *H* (Jianhua *et al*. (1991)):

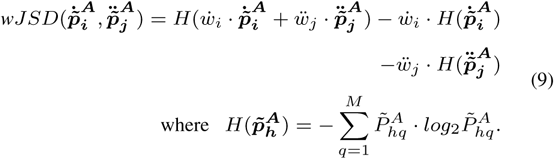

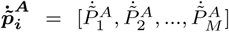, and 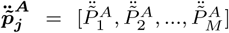 are the corresponding *weighted* discrete probability distributions that describe the probabilities of the reference Atlas genes to appear in the transcriptome of the corresponding tissues. The *weighted* probabilities are calculated as:

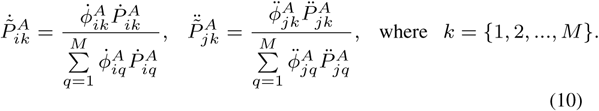

To calculate the *wJSD* (see equation (9)), we need also to determine the weights 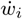 and 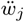 of the corresponding probability distributions 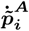 and 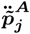. Next, we present a novel method that quantifies our “confidence” on how well the tissue samples 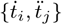 “represent” their corresponding tissue sets 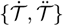, and appropriately adjust the weight values 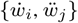.

For each tissue set 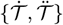, we form the matrices 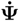 and 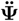 where each of their rows (i.e. 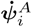 and 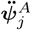) correspond to the standardized versions (e.g. z-scores) of the Atlas signature vectors 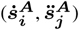 of the corresponding tissue samples.

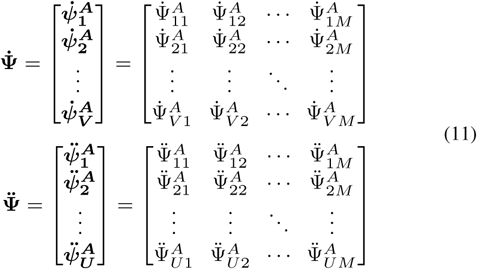

We assume that each standardized reference Atlas signature vector (e.g. 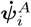 and 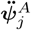) is a random realization of a multivariate normal distribution (e.g. 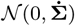 and 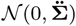). To estimate the covariance matrices 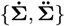 of these distributions we apply the *Graphical Lasso (GL)* algorithm to the corresponding data matrices 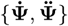. *GL* is a computationally efficient algorithm which has been extensively used to identify gene-gene interaction networks from RNA-seq datasets (Friedman *et al*. (2008)). Its main advantage, is its ability to estimate the precision matrix (i.e. **Ω** = **Σ**^-1^) even in cases where the number of samples is far less than the number of variables (*n<<p*) which holds for transcriptomic datasets (i.e. number of samples *<<* number of genes). In such cases, other covariance estimation methods, such as Maximum Likelihood Estimation, fail since the sample (or empirical) covariance matrix ***𝒮*** is rank deficient. *GL* addresses this limitation using the assumption that precision matrix **Ω** = **Σ**^-1^ is sparse (i.e. the graph structure of the variables’ interactions is sparse). To estimate the precision matrix **Ω**, *GL* efficiently solves the following optimization problem:

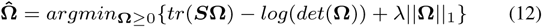

where ‖.‖_1_ is the *L*_1_-norm (i.e. the sum of the absolute values of the elements of Ω); *det*(Ω) is the determinant of Ω; *λ* is the sparsity parameter that controls the density (i.e. the number of edges) of the graphical model; and ***S*** the *M* × *M* sample covariance matrix calculated as ***S*** = **Ψ**^*T*^ **Ψ**. To optimally choose the value of *λ* we use the *StARS* method (Liu *et al*. (2010)) which is stability-based approach for selecting the regularization parameter in high dimensional graphical models.

Using the estimated covariance matrices of the multivariate normal distributions 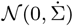 and 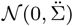 we calculate the likelihood of the tissue samples 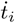 and 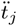 to belong to the corresponding distributions as:

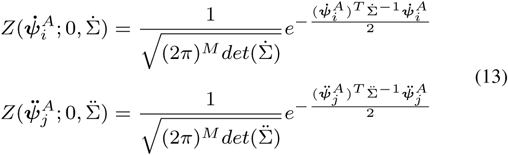

The likelihoods 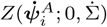 and 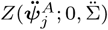, provide information on how “confident” we are that the tissue samples 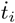 and 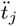 are “good” representatives of the corresponding tissue set 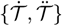. Using this information we calculate the weights 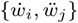 as:

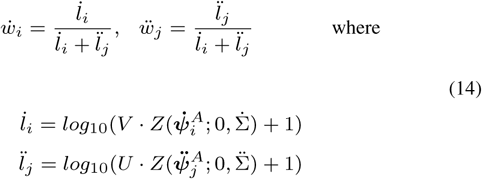

where *V* and *U* are the number of samples in tissue sets 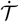 and 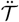 respectively. Note that 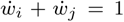. Note that for the calculation of the weight we also consider the sample sizes of the corresponding tissue sets. More specifically, the larger the sample size the larger the weight we assign to the corresponding distribution. This can be justified if we consider that larger the sample sizes provide more confidence about the accuracy of the estimated parameters (i.e. covariance matrix) of the multivariate normal distribution.

#### 2.3.2 The weighted Rankings Correlation Coefficient

The second term of the *TSD* (see equation (4)) is the rankings correlation distance (*RCD*) between the reference Atlas genes ranking vectors 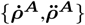 of the compared tissues 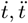. For the case where we have sets of tissue samples (i.e. 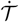 and 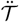) we use *weighted Rankings Correlation Coefficient (wRCC)* which can be calculated as:

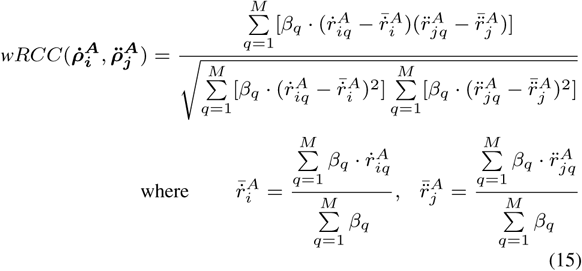

are the corresponding weighted means of the rankings of the Atlas genes. in equation (15) *β*_*q*_, where *q* = {1, …, *M*}, are the weights which assign different “confidence” to the corresponding rankings of the *M* Atlas genes. To calculate the values of these weights, we propose the following method.

Using the ranking vectors 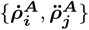 of the Atlas genes of the tissue samples that contained in each set (i.e. 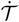 and 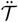) we form the following matrices:

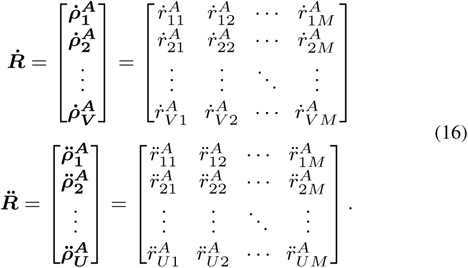

Each row of matrices 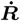 and 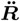 contains the rankings of the expressions of the Atlas genes in the transcriptome of the corresponding tissue sample, and each column includes the rankings of the expressions of a specific Atlas gene across the tissue samples of the corresponding set.

In matrices 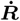 and 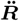, we assume that the rankings of each Atlas gene (e.g. *k*^*th*^ gene) across tissues, were generated by a normal distribution (e.g. 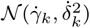 and 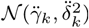) with parameters:

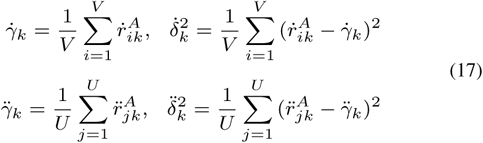

Using this assumption, the likelihood about the rankings of the *k*^*th*^ Atlas gene can be calculated as:

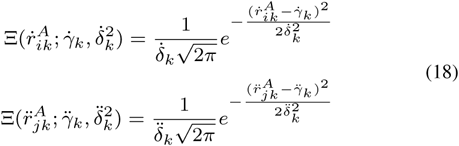

The larger the 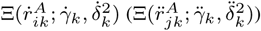 the more “confident” we are about the ranking 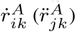 of the *k*^*th*^ Atlas gene, We quantify our “confidence” as:

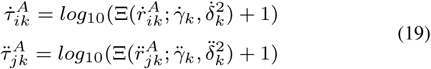

To avoid negative “confidence” values we added “1” before taking the logarithm of the likelihoods. Using the 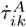 and 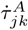 we calculate the importance weight of the *k*^*th*^ Atlas gene as:

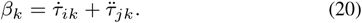

By applying these weights to equation (15) we can calculate the *wRCC* which also takes values in range [-1, 1].

#### 2.3.3 The weighted Transcriptomic Signature Distance

After calculating *wJSD* and *wRCC* we can calculate the weighted version of the transcriptomic signature distance (*wTSD*) as:

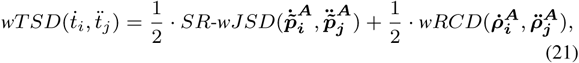

where 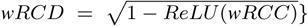 is the *weighted Rankings Correlation Distance*. For the calculation of the *wRCD* we assume that if two vectors have *wRCC <* 0 (i.e. anti-correlated) then their Rankings Correlation Distance is maximum (i.e. 1). Similar to the *TSD* (see equation (4)), *wTSD* takes its values in [0, 1].

## 3 Results and Discussion

We present here extensive results generated using publicly available RNA-seq datasets demonstrating the validity and value of the proposed Transcriptomic Signature Distance (i) for measuring the distance between different organ tissues, (ii) for assessing the distance between healthy and diseased organ tissues, and (iii) for evaluating the similarity of organ models (e.g., organoids, animals, etc.) to the corresponding human organ.

### 3.1 Using *wTSD* with real data to assess distance of human organ tissues

In Section 2.2 we used two hypothetical scenarios to illustrate that using *SR-JSD* or *RCC* alone may fail to adequately capture the transcriptomic differences of tissue samples. In this section, we use real data to show this ineffectiveness and justify the advantages of the proposed *wTSD* as a higher resolution method. For this purpose, we used the publicly available dataset in GEO GSE120795, also presented in Suntsova *et al*. (2019), a comprehensive gene expression database of normal human tissues based on uniformly screened RNA-seq data. This database includes 142 tissue samples taken from 20 organs of healthy human donors of different ages, collected no later than 36 hours after death. The Human Protein Atlas (HPA) includes information for only 13 out of the 20 organs in the database, and these were used in our analysis. As we can see in Table S1 in Supplementary Material the number of samples as well as the number of signature genes identified by the HPA project for each one of the 13 organs used vary considerably. The *wTSD* distance is designed to deal with this kind of situations in a principled manner.

The Heatmaps of Figure 3 depict the mean inter/intra-organ distances. Each row corresponds to a specific organ being used as reference whose HPA signature genes were utilized to estimate the distances of the other organs (columns) from it. As expected, the mean intra-organ tissue sample distances (main diagonal elements) are smaller than the corresponding inter-organ distances (off-diagonal elements of the same row).

**Fig. 3.**
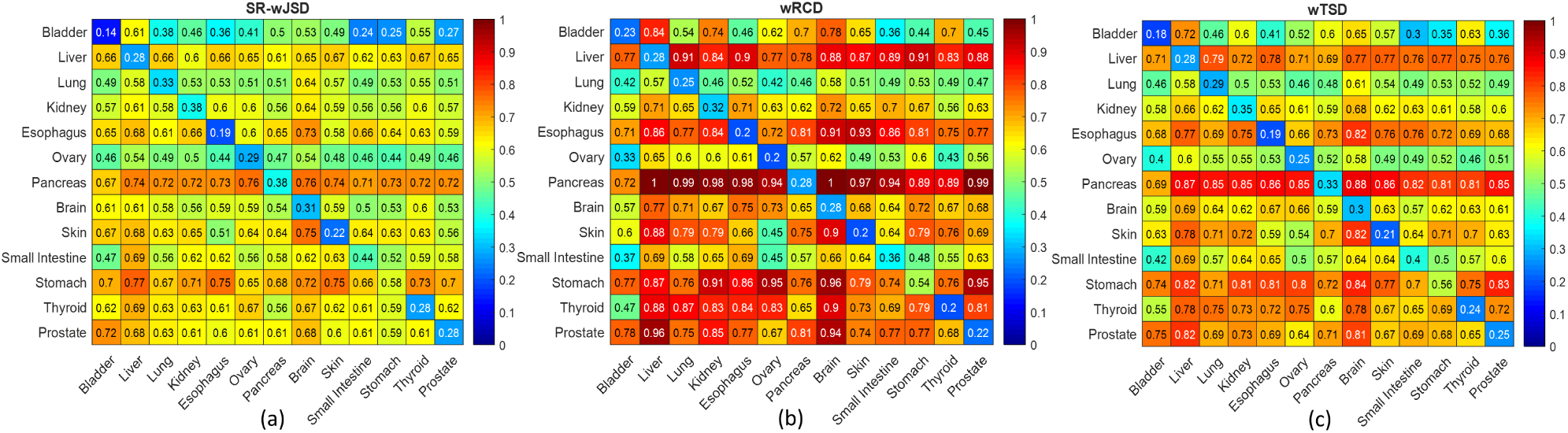
Heatmaps of means of pairwise distances between organ tissue samples. (a) SR-wJSD, (b) wRCD, (c) wTSD. The rows correspond to the reference organs whose Atlas genes were used in the distance calculations.

If we examine carefully corresponding rows in Figures 3a and 3b we see that in many cases there is no correlation between the corresponding *SR-wJSD* and *wRCD* values, indicating that these two pieces of information capture different aspects of transcriptome dissimilarities. To better illustrate this fact, Figures 4a-4b and 4c-4d depict the distances of organs from Lung and Kidney (references) respectively in a 2D-space where *SR-wJSD* and *wRCD* are used as coordinates. In these plots, each organ’s name label is centered at the mean value of the corresponding pairwise tissue sample distances (*SR-wJSD* and *wRCD*) from the reference organ. In the zoomed-in version of Figure 4b we see that the {Liver, Brain} and {Small Intestine, Thyroid} pairs have almost equal *wRCDs* but different *SR-wJSDs* coordinate values. On the other hand, in the zoomed-in Figure 4d, {Thyroid, Esophagus}, {Pancreas, Small Intestine} as well as {Bladder and Prostate} pairs have almost equal *SR-wJSDs* but different *wRCDs* coordinate values. Based on these observations it is clear that the proposed new distance, *wTSD*, which combines the *SR-wJSDs* and *wRCDs* information while also taking into account the intra-class tissue samples variability of every organ (see Section 2), provides a higher resolution picture of the reality.

**Fig. 4.**
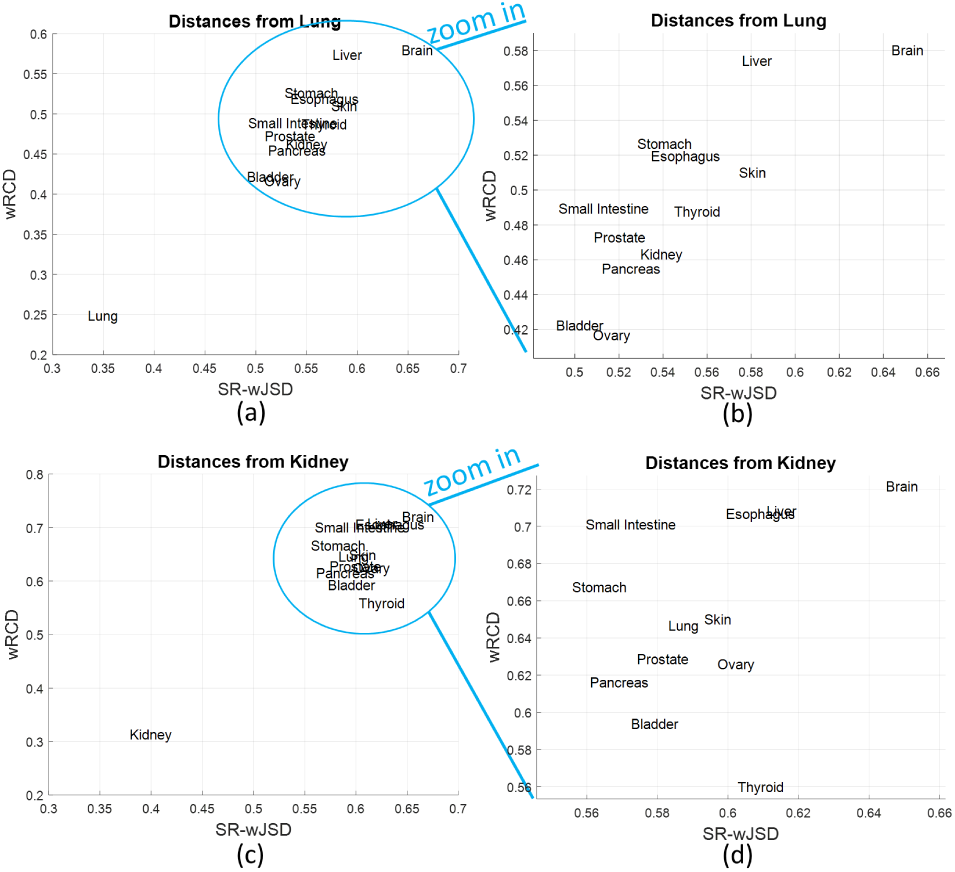
The inter-organ distances from: (a)-(b) Lung and (c)-(d) Kidney used as reference organ. The name label of each organ is centered at the mean value of pairwise distances (SR-wJSD and wRCD) between that organ’s tissue samples and the samples of the reference organ (Lung or Kidney respectively).

### 3.2 Using *wTSD* to assess tissue distance of disease subtypes and progression stages

In this section, we present results demonstrating that *TSD* can be used to resolve transcriptomic distance of normal tissues from tissues of disease subtypes as well as tissues of different disease progression stages. For this purpose, we are using publicly available RNA-seq datasets characterizing two different diseases: idiopathic pulmonary fibrosis and liver cancer.

#### 3.2.1 Using *wTSD* with idiopathic pulmonary fibrosis dataset

Recently published research (McDonough *et al*. (2019)) has studied the progression mechanisms of Idiopathic Pulmonary Fibrosis (IPF), a lethal chronic lung disease which progresses the fibrosis in lungs over time, causing serious breathing difficulties. IPF affects 13 to 20 per 100K people worldwide. According to the National Institute of Health, about 30K to 40K patients in the USA are diagnosed with IPF every year. More than 50% of IPF patients die within 3 − 5 years after the initial diagnosis (Kim *et al*. (2006); Lederer *et al*. (2018)). The RNA-seq dataset of this study, available in NCBI’s Gene Expression Omnibus (GEO GSE124685), consists of 84 samples classified in the following four categories: (i) Controls (n=35), (ii) IPF Early (n=19), (iii) IPF Moderate (n=15) and (iv) IPF Severe (n=15). The samples’ categorization was made based on the extent of lung fibrosis, assessed using microCT quantitative imaging and tissue histology (McDonough *et al*. (2019)). Using this dataset and the information of the 239 genes which, according to HPA, can be considered as the transcriptomic signature of the healthy human lung (see Table S1), we calculated the transcriptomic distances (*SR-wJSD, wRCD* and *wTSD*) of all pairs of tissue samples, one sample belonging in the Controls (reference) group and the other in the diseased groups.

Figure 5a shows the transcriptomic distances (*SR-wJSD* and *wRCD*) of the different IPF progression stages from the healthy lung tissues. The label of each IPF progression stage name is centered at the coordinates of the mean value of the pairwise distances of samples in the corresponding IPF progression group and Control group samples (all pair combinations considered). Figure 5b shows boxplots of the corresponding distributions of the pairwise *wTSD* distances. The results indicate that as the severity of IPF increases, the corresponding transcriptomic signature distance from the Control class also increases. This fact demonstrates the interpretability of the proposed distance. Table S2 in Supplementary Material summarizes the results of the two-sample t-test between the corresponding distributions of the pairwise *wTSD* distances (presented in Figure 5b). The decision of the test is equal to 1 *(h=1)* if the test rejects the null hypothesis (that the groups of the distances have equal means and equal but unknown variances) at the 1% significance level. Table S2 results clearly show that *wTSD* can successfully identify the different IPF progression stages based on the HPA lung signature genes. Moreover, the *wTSD* differences are statistically significant for all comparisons between (i) the Control and IPF stages, and (ii) different IPF stages, a fact that demonstrates the ability of *wTSD* to capture the transcriptomic differences of the corresponding categories.

**Fig. 5.**
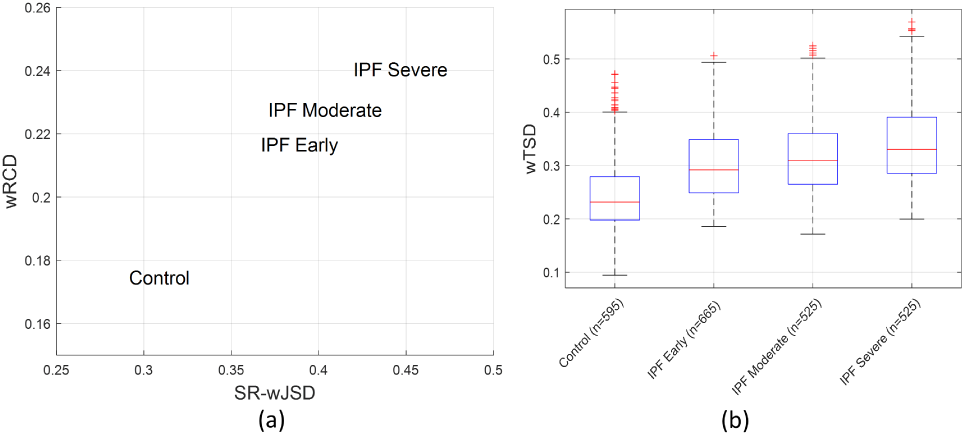
The distances of the different IPF progression stages from the healthy lung. (a) The labels of IPF stages are centered at the coordinates determined by the means of the pairwise distances (SR-wJSD and wRCD) between tissue samples in the corresponding groups and the Controls. (b) Boxplots summarizing the distributions of the corresponding pairwise wTSD distances. Here “n” denotes the number of pairwise distances (wTSD) calculated between the Controls and the samples in the different IPF progression stages.

#### 3.2.2 Using *TSD* with human liver cancer dataset

In a recent study Broutier *et al*. (2017), human primary liver cancer-derived organoids were employed to recapitulate the pathophysiology of human liver tumors. From the provided RNA-seq dataset (GEO GSE84073), we extracted samples for healthy human liver tissue (Controls) and different human liver tumor subtypes, in particular: Hepato-Cellular Carcinoma (HCC), Cholangio-Carcinoma (CC) and combined HCC/CC (CHC). The number of tissue samples in each group was relatively small: (i) Controls (n=4), (ii) HCC (n=3), (iii) CC (n=4) and (iv) CHC (n=3). We also used for every sample the expression information of the 936 genes, which, according to HPA, form a transcriptomic signature of the healthy human liver (see Table S1). Next, using as reference organ the healthy human liver, we computed the corresponding pairwise distances (*SR-JSD, RCD* and *TSD*) between its samples and samples in the different cancer subtype groups. We remark here that due to the limited number of samples in the cancer groups, we decided to use the simple version (not weighted) of the *TSD*.

Figure 6a shows the transcriptomic distances (*SR-JSD* and *RCD*) of the different cancer subtypes from the healthy liver. Each cancer subtype’s name label is centered at the coordinates of the mean value of the pairwise distances between samples in the corresponding cancer group and Control samples. Figure 6b shows boxplots of the distributions of these pairwise *TSD* distances. These results demonstrate the ability of *TSD* to represent the distance difference of controls from the tumor subtypes. It is interesting to remark that the distance of the CHC group (tumor tissue, which is a combination of HCC and CC) from the Controls is in-between the corresponding distances of the HCC and CC, a fact that conforms with our human intuition. Moreover, Table S3 in the Supplementary Material summarizes the results of the two-sample t-test between the corresponding distributions of the pairwise *TSD* distances (presented in Figure 6b). In Table S3, the decision of the test is equal to 1 *(h=1)* if the test rejects the null hypothesis at the 5% significance level. The results show that all comparisons except one (Control vs. HCC) have significantly different *TSD* distances, which indicates the ability of *TSD* to identify the transcriptomic differences of the corresponding groups.

**Fig. 6.**
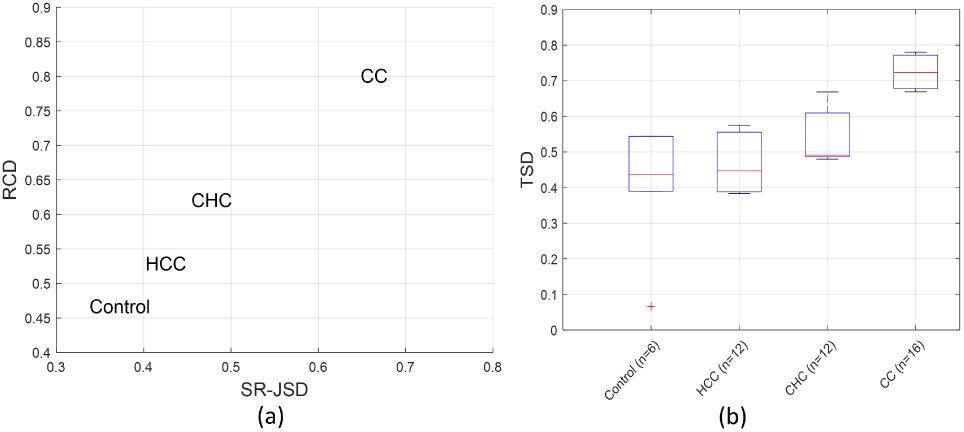
Distances of different liver cancer subtypes stages from the healthy liver. (a) The liver cancer subtype name labels are centered at the coordinates determined by the means of the pairwise distances (SR-wJSD and wRCD) between the samples in the corresponding subtype groups and the Controls. (b) Boxplots summarizing the distributions of the corresponding pairwise TSD distances. Here “n” is the number of pairwise distances (TSD) calculated between Control group tissue samples and different cancer subtype group samples.

### 3.3 Assessing distance of human organs from different organ models

In this section, we show that the proposed transcriptomic signature distance can be used to assess the “physiological relevance” of different organ models to a human organ based on RNA-seq data. Specifically, we computed the transcriptomic signature distance of: (i) mus musculus (mouse) organs, (ii) rattus norvegicus (rat) organs, and (iii) human-derived organoids from the human liver and kidney used as reference organs. We obtained the RNA-seq data for the healthy human organ tissues from the publicly available database developed by Suntsova *et al*. (2019) (described in Section 3.1). The number of available samples for each healthy human organ is provided in Table S1. For mouse and rat, we obtained RNA-seq data from the database developed by Sollner *et al*. (2017). For both species, the number of available samples per organ was equal to 3. Finally, we retrieved RNA-seq data for healthy human liver and kidney derived organoids from the publicly available datasets (GSE84073 and GSE99582) presented in Broutier *et al*. (2017) and Phipson *et al*. (2017) respectively. The number of samples of the healthy liver and kidney organoids was 6 and 3, respectively. To compare the transcriptomic signatures between the different species, we associated the mouse and rat genes to human homologous genes using the R-package *biomart* (Smedley *et al*. (2015)). Due to the limited number of samples (*n* = 3) in some of the categories under comparison, we used the “simple” version of *TSD* (not weighted).

Figures 7a and 8a show, for the liver and kidney datasets respectively, the pairwise distances (*SR-JSD* and *RCD*) of organ model samples (mouse, rat, organoids) from healthy human organ tissue samples. In both Figures, a circle depicts the distance of a model tissue sample (represented by color black, blue, red) to a control sample. On the other hand green circles are used for pairwise distances between samples of the control group. The boxplots in Figures 7b and 8b summarize the distributions of these pairwise distances. The results indicate that the liver tissue samples of mouse and rat are transcriptomically closer to the human liver than the human-derived liver organoid samples. However, for the kidney, the transcriptomic signature distances of the mouse, rat and human derived kidney organoids, from the healthy human kidney, are very similar. Another interesting observation is that for both organs, the transcriptomic distances of mouse and rat from the corresponding healthy human organs are similar. As shown in Sudmant *et al*. (2015) mouse and rat have similar transcriptomic profiles between homologous organ tissues which is also confirmed by Figure 7 and Figure 8. Table S4 and Table S5 summarize for liver and kidney respectively the results of the two-sample t-tests between the corresponding distributions (see Figures 7b and 8b) of the pairwise *TSD* distances.

**Fig. 7.**
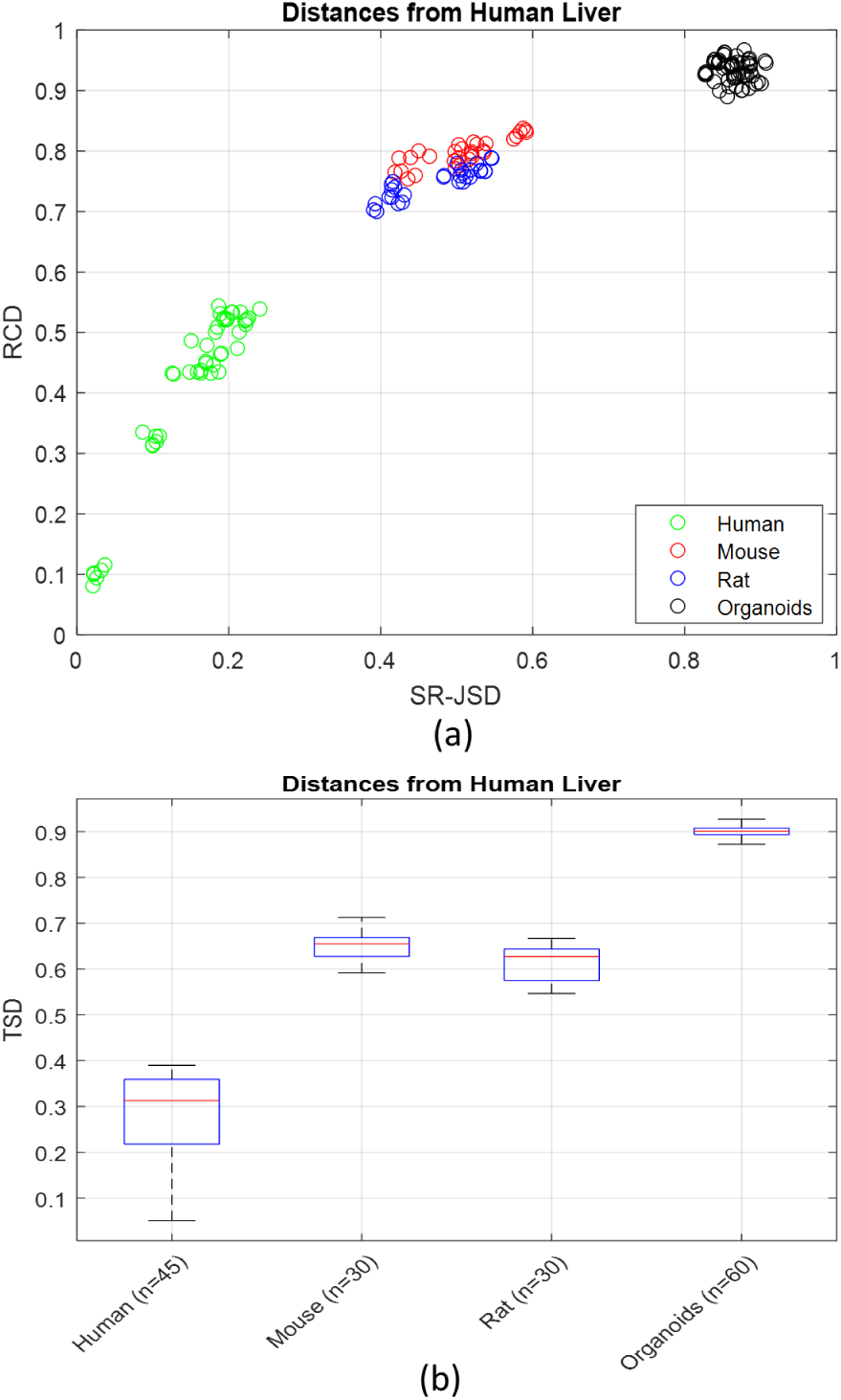
The distances of the different liver models from the human liver. (a) The pairwise distances (SR-JSD and RCD) of organ model tissue samples (mouse (red circles), rat (blue circles), organoids (black circles)) from the corresponding healthy human organ tissue samples. (b) Boxplots summarizing the distributions of corresponding pairwise TSD distances. Here “n” is the number of pairwise distances (TSD) calculated between tissue samples of liver organ models and human liver.

**Fig. 8.**
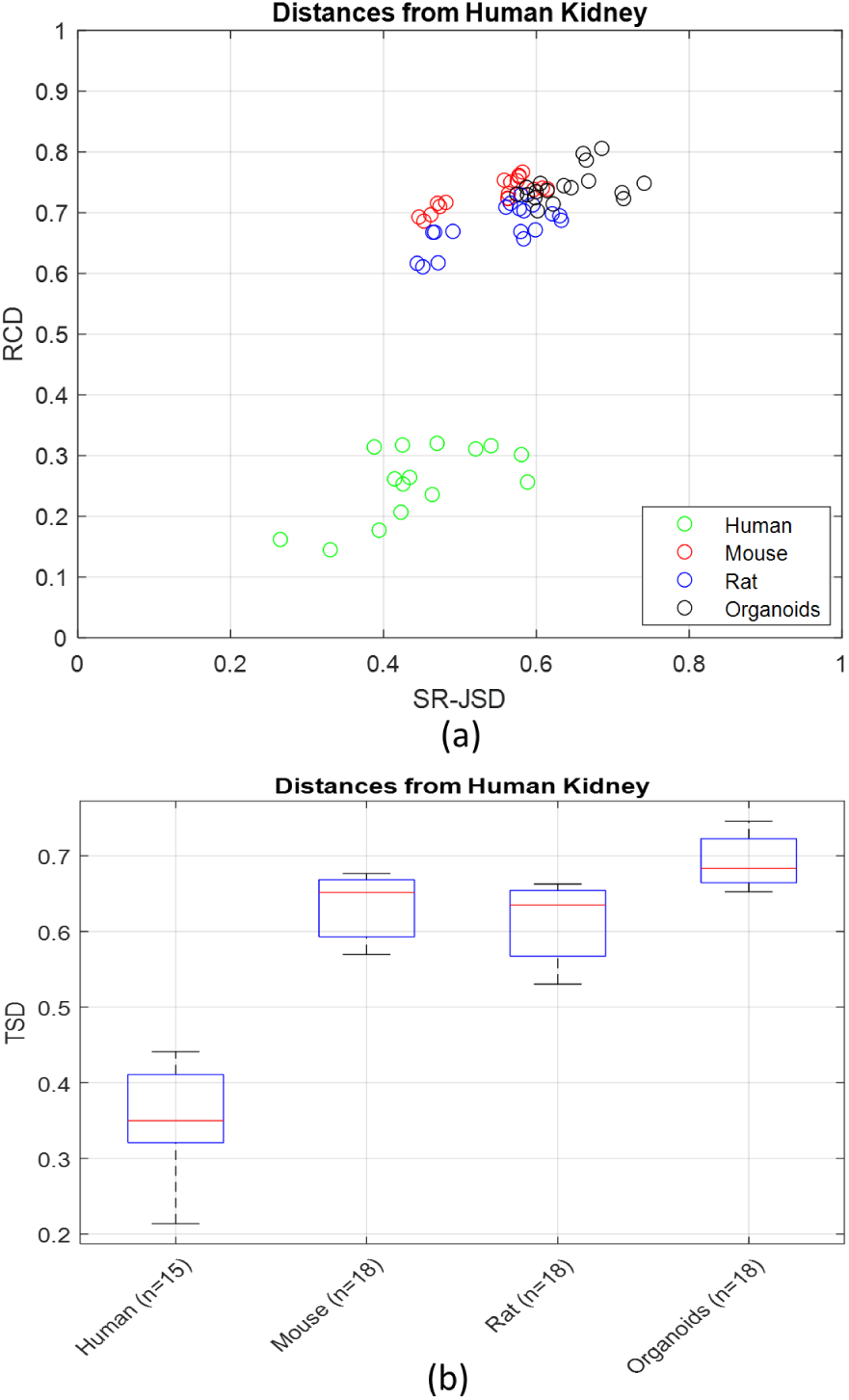
The distances of different kidney models from the human kidney. (a) Pairwise distances (SR-JSD and RCD) of organ model samples represented as small circles; mouse (red), rat (blue), organoids (black) from the corresponding healthy human organ tissue samples. (b) Boxplots are summarizing the distributions of corresponding pairwise TSD distances. Here “n” is the number of pairwise distances between tissue samples of kidney organ models and human kidney tissue samples.

At this point we should mention a very interesting property of the *(w)TSD*. When all the compared tissues (e.g., 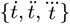) represent the same organ (i.e. are all compared based using the same set of signature genes) as the case here for liver and kidney, *(w)TSD* satisfies all the necessary conditions of a true metric. Based on *SR-(w)JSD* and *(w)RCD* characteristics (Jianhua *et al*. (1991); Jiaxing *et al*. (2019)) it is a matter of simple algebra to show that:

1. Is symmetric: 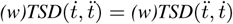
2. Is non-negative: 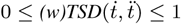
3. 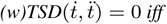: 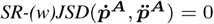 and 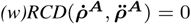
4. 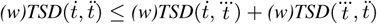 (triangle inequality), ∀ triplet of tissues 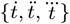.

In this section, we presented how to use *(w)TSD* as a distance to measure the similarity between organ tissue samples in different scenarios arising in practice. We remark that we can also employ *(w)TSD* in a variety of other situations for measuring distance of biological samples as long as we have access to their gene expression data and information about their signature genes. For example, *(w)TSD* can be used to assess the distances between different cell types using single-cell RNA-seq data and information about cell-type-specific signature gene sets that we can retrieve from publicly available databases or the expanding literature on the subject (Kotliar *et al*. (2019); Merienne *et al*. (2019); Thul *et al*. (2017)).

## 4 Conclusions

We presented the development of *TSD*, a new distance we have introduced for quantifying the transcriptomic similarity of organ tissues. The development of *TSD* is grounded on information theory and advanced statistics. Also, *TSD* exploits the availability of “signature” genes for human organs, provided in the well-curated publicly available HPA database, to emphasize organ tissue differences and mask the effects of measurement noise and inter-donor variability in the distance calculations. We also presented a novel method that considers the gene expression and ranking variations across homologous tissue samples and appropriately incorporates this information into the distance calculations. We justified the effectiveness and reliability of the proposed distance and evaluated its performance using many different publicly available RNA-seq datasets. We presented extensive experimental results that validate the ability of *TSD* to represent distances between different organ tissues coherently. Moreover, we have shown how *TSD* can be used to assess the distance of alternative organ model technologies (in-vivo, ex-vivo, etc.) to the corresponding human organ. To the best of our knowledge, *TSD* is the first distance based on information theory, which allows us to assess the similarity of model organ tissue samples to the human organ they represent based on a reference gene set. We are confident that *TSD* can be a valuable tool in many disciplines, such as tissue engineering, microphysiolocal systems design, single-cell types comparison, etc. To facilitate its use we plan to make its R code implementation available on Github.

