## Supplementary Material for "An information-theoretic approach for measuring the distance of organ tissue samples using their transcriptomic signatures"

**Table S1**: Number of Quality Control-passed samples and number of signature genes identified by the Human Protein Atlas (HPA) for each one of the 13 healthy tissues used (dataset by Suntsova et al. (2019))

| **Organs Suntsova et al. (2019)** | **Number of QC-passed samples Suntsova et al. (2019)** | **Number of signature genes in HPA** |
|---|---|---|
| Bladder | 4 | 99 |
| Liver | 10 | 936 |
| Lung | 7 | 239 |
| Kidney | 6 | 413 |
| Esophagus | 10 | 311 |
| Ovary | 4 | 173 |
| Pancreas | 6 | 422 |
| Brain | 7 | 488* |
| Skin | 6 | 547 |
| Small Intestine | 5 | 764 |
| Stomach | 10 | 159 |
| Thyroid | 6 | 199 |
| Prostate | 6 | 120 |

*Note that for the Brain we used as signature genes the stricter HPA set of *enriched genes*. This allowed us to reduce the number of signature genes from 2587 to 488 and significantly cut down the execution time of the Graphical Lasso algorithm.

**Note:** In the following tables the decision of the *t-test* is equal to *1 (h=1)* if the test rejects the *null hypothesis* (i.e. the groups of the compared *TSD* distances have equal means and equal but unknown variances) at the 5% significance level.

**Table S2**: Results of the two-sample *t-test* between the w*TSD* distributions (see boxplots in Figure 5b) of the Control and IPF progression stages.

| **Comparison** | ***h*** | ***p-value*** |
|---|---|---|
| Control vs. IPF Early | 1 | $2.62*10^{-53}$ |
| Control vs. IPF Moderate | 1 | $3.81*10^{-64}$ |
| Control vs. IPF Severe | 1 | $1.41*10^{-100}$ |
| IPF Early vs. IPF Moderate | 1 | 0.0013 |
| IPF Early vs. IPF Severe | 1 | $1.2*10^{-21}$ |
| IPF Moderate vs. IPF Severe | 1 | $1.09*10^{-9}$ |

**Table S3**: Results of the two-sample *t-test* between the *TSD* distributions (see boxplots in Figure 6b) of the Control and liver cancer subtypes.

| **Comparison** | ***h*** | ***p-value*** |
|---|---|---|
| Control vs. HCC | 0 | 0.31 |
| Control vs. CHC | 1 | 0.04 |
| Control vs. CC | 1 | $1.09*10^{-6}$ |
| HCC vs. CHC | 1 | 0.04 |
| HCC vs. CC | 1 | $1.19*10^{-10}$ |
| CHC vs. CC | 1 | $2.96*10^{-8}$ |

**Table S4:** Results of the two-sample *t-test* between the *TSD* distributions (see boxplots in Figure 7b) of the Human Liver and the Liver emulation models (in-vivo and in-vitro).

| **Comparison** | ***h*** | ***p-value*** |
|---|---|---|
| Human Liver vs. Mouse Liver | 1 | $3.32*10^{-30}$ |
| Human Liver vs. Rat Liver | 1 | $5.23*10^{-27}$ |
| Human Liver vs. Hum. Liver Organoids | 1 | $3.43*10^{-71}$ |
| Mouse Liver vs. Rat Liver | 1 | $8.61*10^{-5}$ |
| Mouse Liver vs. Hum. Liver Organoids | 1 | $2.21*10^{-64}$ |
| Rat Liver vs. Hum. Liver Organoids | 1 | $5.39*10^{-68}$ |

**Table S5**: Results of the two sample *t-test* between the *TSD* distributions (see boxplots in Figure 8b) of the Human Kidney and the Kidney emulation models (in-vivo and in-vitro).

| **Comparison** | ***h*** | ***p-value*** |
|---|---|---|
| Human Kidney vs. Mouse Kidney | 1 | $7.19*10^{-16}$ |
| Human Kidney vs. Rat Kidney | 1 | $2.79*10^{-14}$ |
| Human Kidney vs. Hum. Kidney Organoids | 1 | $1.06*10^{-18}$ |
| Mouse Kidney vs. Rat Kidney | 0 | 0.14 |
| Mouse Kidney vs. Hum. Kidney Organoids | 1 | $4.75*10^{-5}$ |
| Rat Kidney vs. Hum. Kidney Organoids | 1 | $1.52*10^{-6}$ |
